# Deregulation of VEGFR-2 and PGFR expression and microvascular density in a triple-negative model of canine malignant mammary tumors with lymph node or lung metastasis

**DOI:** 10.1101/490144

**Authors:** Denner Santos dos Anjos, Aline Fernandes Vital, Patrícia de Faria Lainetti, Antonio Fernando Leis-Filho, Fabiola Dalmolin, Fabiana Elias, Sabryna Gouveia Calazans, Carlos Eduardo Fonseca-Alves

**Affiliations:** Veterinary Science Graduate Program, University of Franca (UNIFRAN), Franca, Brazil; Department of Veterinary Surgery and Anesthesiology, School of Veterinary Medicine and Animal Science, São Paulo State University – UNESP, Botucatu, SP, Brazil; Federal University of the Southern Border, Realeza, PR, Brazil

**Keywords:** angiogenesis, metastasis, mammary neoplasm

## Abstract

Canine mammary tumors (CMT) are the most common cancer in noncastrated female dogs. Interestingly, triple-negative tumors are the most common molecular subtype in female dogs. In this study, we proposed to evaluate the expression of VEGFR-2, PDGFR and microvascular density (MVD) in a group of metastatic and nonmetastatic triple-negative CMT and compare the expression based on clinical parameters. Twenty-six female dogs with triple-negative mammary tumors were divided into three groups: nonmetastatic tumors (NMT) (N=11), tumors with lymph node metastasis (LNM) (N=10) and tumors with lung metastasis (LM) (N=5). We observed increased VEGFR-2 expression in LNM compared with NMT and a positive correlation between tumor grade and VEGFR-2 expression. A positive correlation was noted between VEGFR-2 and PDGFR expression. Regarding microvascular density (MVD), we identified a higher number of vessels in primary tumors with lymph node metastasis and lung metastasis compared with tumors with no metastasis. The primary tumors with lung metastasis exhibited an increased MVD compared with carcinoma with lymph node metastasis. Overall, our results suggest a deregulation of VEGFR-2 and PDGFR and high MVD in metastatic tumors, indicating a role for angiogenesis in tumor progression.

## 1. Introduction

Canine mammary tumors (CMT) are the most common tumor in noncastrated female dogs with a variable clinical behavior [1]. The incidence rates for CMT depends on the geographic origin given that it is a tumor with higher prevalence in countries where castration is not routinely performed [2]. In Brazil, the prevalence of CMT in intact female dogs is approximately 28% to 45% of all tumors in dogs [3, 4]. CMTs resemble human breast cancer (BC), and dogs represent an interesting model for comparative studies. The recent GLOBOCAN estimates of cancer presented an expectation of 2,0938,76 new cases of BC worldwide and 626, 679 deaths related to BC [5].

BC is the most important tumor in women as it is the most diagnosed cancer and the second cause of death related to cancer [6]. Human BC is subdivided into molecular subtypes, such as HER2 enriched, Luminal A, Luminal B and basal-like [6]. Triple-negative tumors are very important as these tumors represent a therapeutic challenging, and limited therapeutic options are available compared with other subtypes [7]. Recently, a study evaluating a large number of cases subdivided CMT into molecular subtypes and found an increased prevalence of triple-negative tumors in dogs [8]. These results indicate that female dogs serve as a natural model for human BC.

In humans, vascular endothelial growth factor-A (VEGF-A) expression increases based on tumor grade. Thus, a tumor with a higher histological grade presents higher VEGF-A levels. Moreover, increased VEGF-A expression correlates with tumor metastasis, indicating the role of the VEGF pathway in human tumors. Vascular endothelial growth factor receptor 2 (VEGFR-2) is one of the principal mediators of VEGF-A activity [9]. VEGF-A and VEGFR-2 levels are associated with the worst outcome in patients with BC. Thus, the VEGF-A/VEGFR-2 signaling pathway exhibits prognostic and predictive value in female BC [9]. VEGF expression was previously investigated in CMT. A correlation between VEGF expression and tumor angiogenesis was observed [10], and VEGF overexpression correlates with lymph node metastasis [11]. However, information regarding VEGFR-2 expression in CMT in the literature is lacking [12].

PDGFR and c-KIT expression is widely studied in human oncology, and both markers exhibit predictive value in human BC. Imatinib mesylate (Gleevec^®^) was previously evaluated in advanced/metastatic breast cancer expressing c-KIT or PDGFR [13, 14]. However, both studies evaluated a low number of patients due to imatinib toxicity. Thus, it was concluded that imatinib mesylate as a monotherapy does not provide a clinical benefit for BC-affected patients and is associated with important side effects [13, 14]. However, PDGFR and c-KIT is overexpressed in human BC and still represent important predictive markers. New studies evaluating other PDGFR/c-KIT inhibitors represent a new therapeutic perspective.

In dogs with mammary tumors, c-KIT exhibits a controversial role in tumorigenesis [15, 16, 17]. In general, c-KIT is expressed in normal mammary glands. During cancer progression, tumor cells lack c-KIT expression [15, 16, 17]. Regarding PDGFR, one previous study evaluated gene expression in CMT [17], and no previous study demonstrated PDGFR expression in CMT. The toxicity of different tyrosine kinase inhibitors has been studied in dogs [18, 19, 20]. Thus, dogs represent an important preclinical model for human cancers.

In humans and dogs, the development of metastasis is the major cause of cancer-related deaths [6, 15]. The PDGFR and PDGF signaling pathway is responsible for intratumoral lymphogenesis, promoting nodal metastasis [21], and the VEGF/VEGFR pathway induces neovasculogenesis [Shibuya, 2011]. Microvascular density (MVD) is very important for tumor progression and is induced by the production of proangiogenic factors by tumor cells [23]. MVD in BC is correlated with overall survival and disease-free interval in both humans [23, 24] and dogs [25]. Given the importance of dogs as a natural model for human BC, this research aimed to evaluate VEGF and PDGFR expression and assess MVD in metastatic and nonmetastatic CMT.

## 2. Material and methods

### 2.1 Study design

This was a prospective nonrandomized study including 26 female dogs from three institutions: Veterinary Teaching Hospital of University of Franca (UNIFRAN), the Veterinary Teaching Hospital of São Paulo State University (UNESP) and the Veterinary Teaching Hospital of the Federal University of the Southern Border (UFFS). All procedures were performed in accordance with the national and international guidelines for use of animals in research. This study was approved by each institutional Ethics Committee in the Use of Animals. The tissue samples were collected between May 2015 and September 2017.

We exclusively included patients with malignant tumors meeting the following criteria: a sufficient amount of tissue in the primary tumor and metastatic foci for immunohistochemical evaluation, received no previous systemic treatment, at least one year of clinical follow-up, only one tumor in the mammary gland and sentinel lymph node evaluation.

### 2.2 Patients

We included female dogs with only one mammary gland lesion, independent of the tumor size or location. All dogs underwent a previous cytological examination indicating a mammary gland tumor. The patients underwent sentinel lymph node assessment according to Beserra et al. [26]. Then, unilateral chain mastectomy was performed. The surgical specimens were stored in 10% buffered formalin for 24 hours. Then, histological processing was performed. Briefly, 4-μm tissue sections were processed for hematoxylin and eosin staining. The histological classification was performed according to Goldschmidt et al. [27], and tumor grade was evaluated according to Karayannopoulou et al. [28].

### 2.3 Clinical evaluation

All patients underwent three-view thoracic radiographic examination, abdominal ultrasound and complete blood count. Then, we obtained the clinical stage from the staging system established by the World Health Organization for CMT and modified by Sorenmo et al. [2]. Patients were classified as stage I-V [2]. Patients with at least stage III disease and at least tumor grade II received adjuvant chemotherapy with four cycles of 300 mg/m^2^ carboplatin and 5 mg/kg of firocoxib every 4 hours for six months Bonolo et al. [29]. Clinical follow-up was performed every three months in the first year with three-view thoracic radiographic examinations and complete blood counts.

### 2.3 Molecular phenotype

We exclusively included patients with triple-negative mammary tumors. Immunohistochemistry was performed according to Abadie [8]. Then, we used a combination of ERα, PR, HER-2, Ki67, CK5/6 and EGFR to classify the different molecular phenotypes (Figure 1).

**Figure 1.**
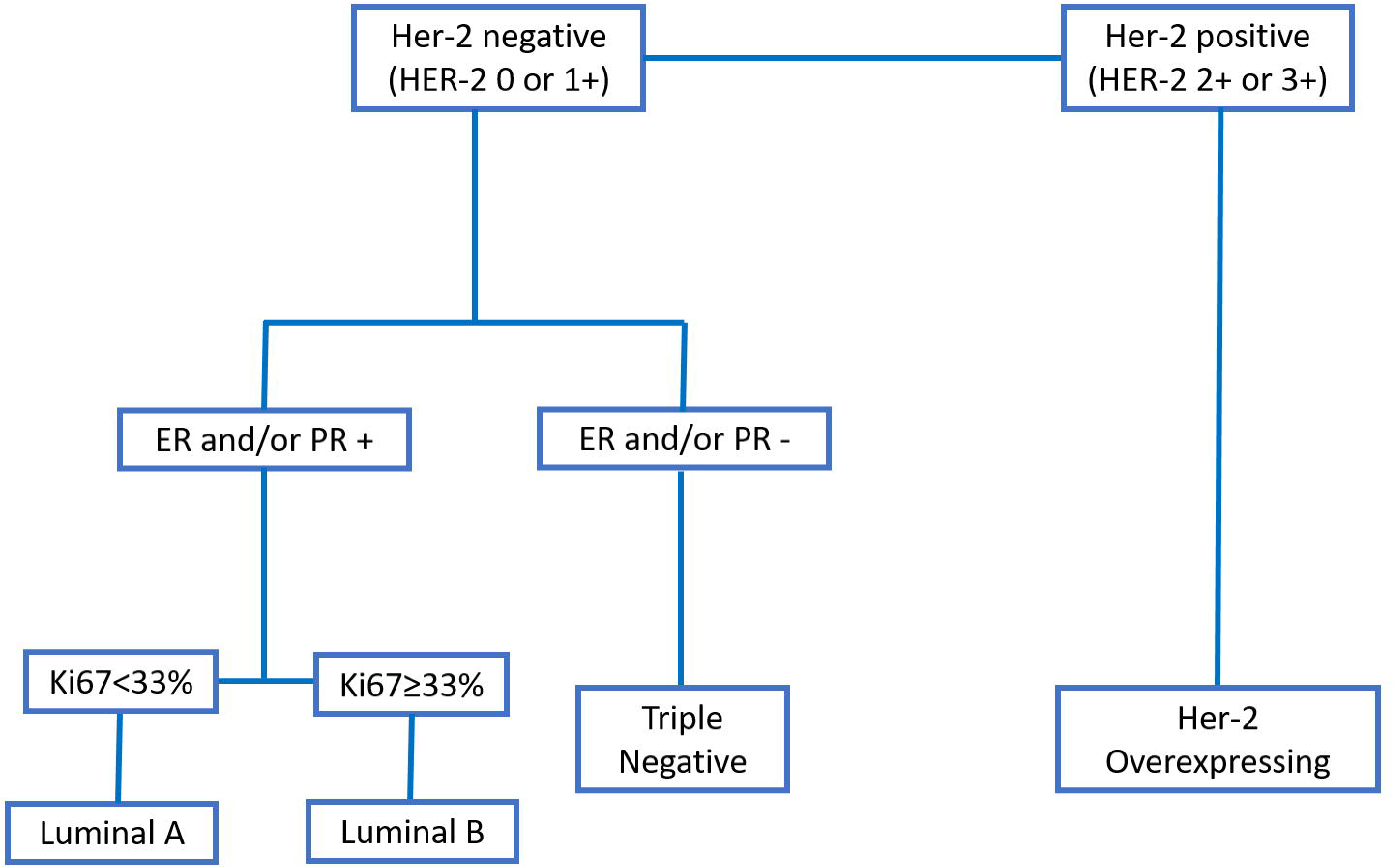
Classification of the molecular phenotypes of canine mammary carcinomas according to each immunohistochemical marker.

### 2.4 Tumor groups

Twenty-six patients met our inclusion criteria, and four samples were used. Our patients were divided into three groups: patients with malignant mammary tumor with no local or distant metastasis and at least one year of follow-up of evaluation (G1, nonmetastatic tumors), patients with malignant mammary tumor with metastasis to sentinel lymph nodes at diagnosis and no distant metastasis (G2, tumors with lymph node metastasis) and patients with malignant mammary tumors with negative sentinel lymph nodes at diagnosis and developed late lung metastasis (G3, tumors with distant metastasis).

### 2.4 Immunohistochemistry

Immunohistochemical evaluation was performed in 41 paraffin blocks: 11 primary tumors from G1, 10 primary tumors and 10 lymph node metastasis from G2 and five primary tumors and five lung metastasis from G3. Charged slides with 4-μm tissue sections were cut, deparaffinized and submitted to antigen retrieval with citrate buffer pH 6.0 in a pressure cooker (Pascal, Dako, Carpinteria, CA, USA). Endogenous peroxidase was blocked with 8% hydrogen peroxide diluted in methanol for 10 minutes. Then, monoclonal VEGFR (Flk1, Abcam, Cambridge, UK), rabbit monoclonal PGFR-β (clone 26E1, Cell signaling, Danvers, MA, USA) and rabbit polyclonal CD31 (ThermoFisher Scientific, Waltham, MA, EUA) antibodies at 1:300, 1:200 and 1:50, respectively, were applied for 18 hours. Afterward, incubation with secondary antibody (Envision, Dako, Carpinteria, CA, USA) for 1 hour was performed, and samples were incubated with 3,3’-diaminobenzidine (DAB, Dako, Carpinteria, CA, USA) for 5 minutes. Counterstaining was performed with Harris hematoxylin for 1 minute. The positive controls were selected according to the Protein Atlas recommendations (https://www.proteinatlas.org). For VEGFR, canine liver was used as a positive control. For PGFR-β, normal testis was used as a positive control. For CD31, we used an internal control (blood vessel in each tumor sample). Mouse (Negative Control Mouse, Dako, Carpinteria, CA, USA) and rabbit immunoglobulin (Negative Control Rabbit, Dako, Carpinteria, CA, USA) were used as negative controls. VEGFR, PGFR-β and CD31 cross-reactivities with canine tissue were provided by the manufacturer.

For VEGFR and PGFR-β, the samples were evaluated by optical microscopy using a semiquantitative score of 0 to 4 [30]. Briefly, 0: absence of labeling, 1: 1% up to 25% of positive cells, 2: 26% up to 50% positive cells, 3: 51 up to 75% positive cells and 4: > 75% positive cells. For Factor CD31, microvessel counts were performed in five fields using the 20x objective, and the mean of the sum of the five fields was used according to Weidner [23].

### 2.5 Statistical evaluation

The results were previously submitted to Shapiro-Wilk normality tests and analysis of variance (ANOVA). If the variables presented a Gaussian distribution, Tukey’s test or nonparametric Kruskal-Wallis test was used for microvascular density analysis. Spearman’s test was used to investigate correlations between variables. Regarding the VEGFR and PDGFR immunoexpression, Chi-square or Fisher exact tests were performed. Statistical analyses were performed using the GraphPad Prism^®^ program (version 6.0 - GraphPad Software, Inc. 2015) with a significance level of 0.05.

## 3. Results

### 3.1 Clinical and Pathological evaluation

Regarding the pathological parameters, tubulopapillary carcinoma was the most commonly observed diagnosis (8/26) followed by solid carcinoma (7/26), complex carcinoma (5/26), comedocarcinoma (4/26) and mixed carcinoma (2/26). Seven carcinomas were classified as grade 1, eight as grade 2 and eleven as grade 3 (Table 1). The clinical parameters are described in Table 1. The mean survival time for all patients independent of the metastatic status was 384.96 days (±123.6) (Figure 2A). Patients with nonmetastatic disease at the diagnosis experienced an increased survival time compared with patients with lymph node metastasis (P=0.0359) (Figure 2B). Patients with grade III tumors experienced a shorter survival time compared with grades I and II (P=0.534) (Figure 2C). A negative correlation was observed between tumor grade and overall survival (P=0.0274; Spearman R= -0.4244). Thus, patients with high tumor grade experienced a reduced survival time (Figure 2D).

**Table 1.**
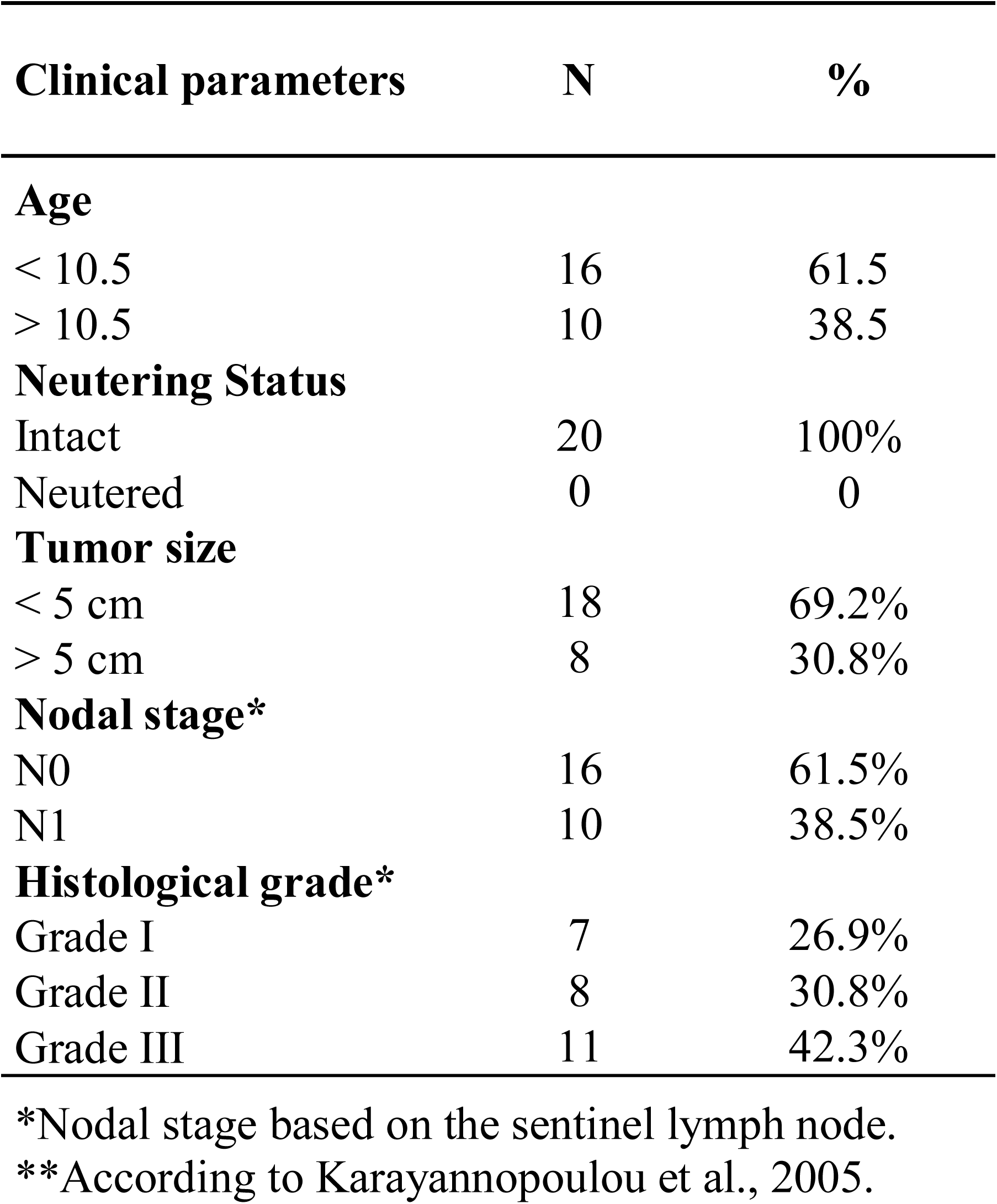
Clinical parameters of female dogs affected by mammary gland tumors.

**Figure 2.**
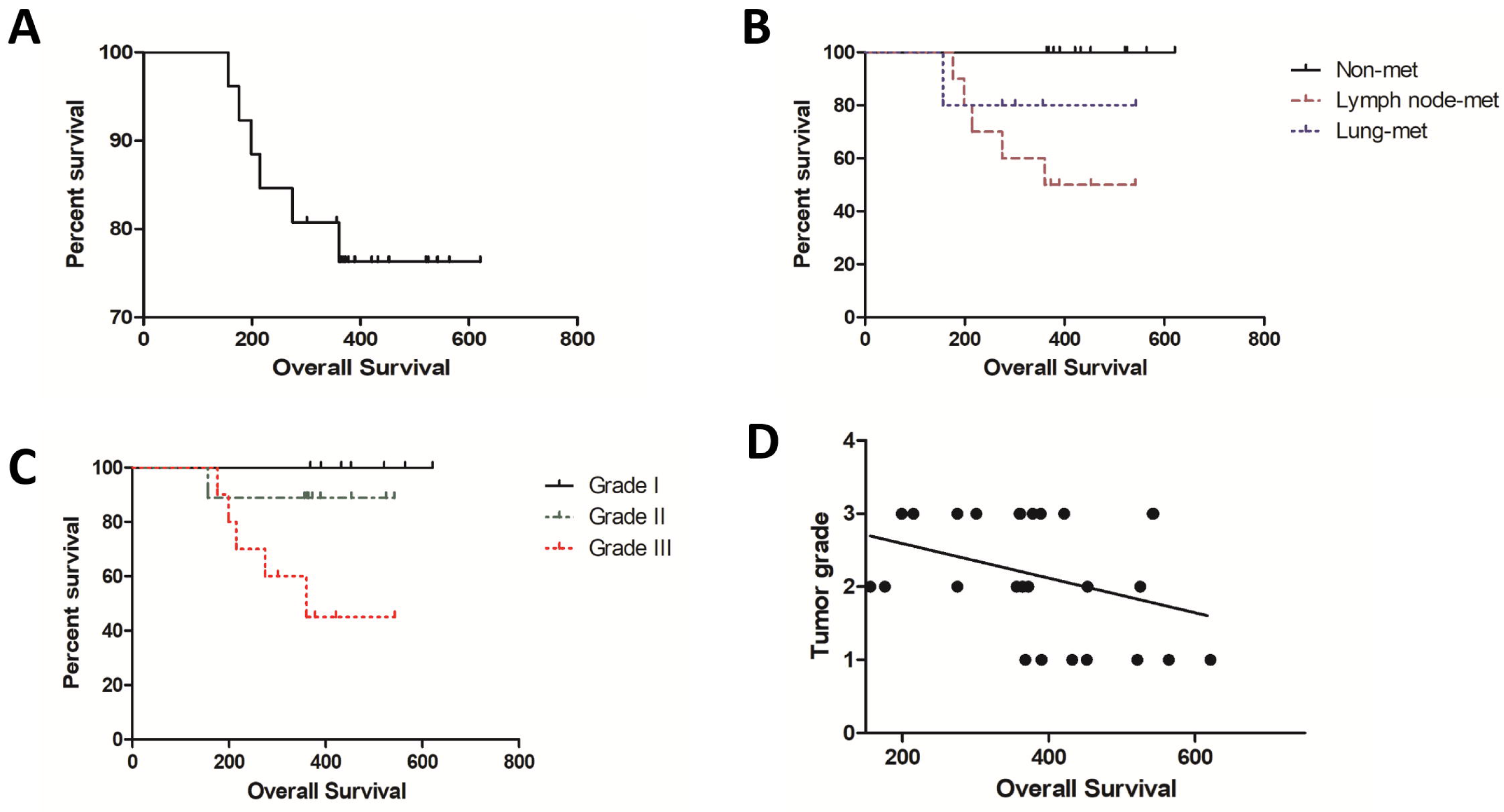
Overall survival of dogs with mammary carcinomas based on clinical parameters. A: Percent survival of all female dogs independent of metastasis status. B: Female dogs with nonmetastatic tumors exhibited increased survival time followed by patients with lung metastasis and lymph node metastasis. C: Overall survival independent of metastasis status according to tumor grade. Patients with grade III experienced a reduced survival time. D: Negative correlation between tumor grade and overall survival. Patients with low-grade tumors exhibited increased survival time.

### 2.2 Immunohistochemistry

We identified VEGFR-positive expression in all primary and metastatic samples. Patients exhibiting lymph node metastasis at diagnosis exhibited increased VEGFR expression compared with nonmetastatic carcinomas (P=0.0238). On the other hand, we did not observe a significant difference when we compared primary carcinomas with lung metastasis with nonmetastatic carcinomas (P=0.1239). We did not observe a significant difference between lymph node metastasis and lung metastasis (P=0.7243). We also did not observe a significant difference when comparing the primary carcinomas with its respective metastasis. We identified a positive correlation between tumor grade and VEGFR expression (P=0.001; Spearman R=0.6071). No correlation between VEGFR expression and overall survival was observed (P=0.125; Spearman R=-0.3087). VEGFR immunoexpression results are presented in Figure 3 and Table 2.

**Table 2.** Immunohistochemical evaluation of VEGFR-2, PDGFR and microvascular density in canine mammary gland tumor samples.

|  | VEGFR-2 score |  |  |  |  | PDGFR score |  |  |  |  | Microvascular Density |
| --- | --- | --- | --- | --- | --- | --- | --- | --- | --- | --- | --- |
|  | 0 | 1 | 2 | 3 | 4 | 0 | 1 | 2 | 3 | 4 | Mean (SD*) |
| **Non-metastatic Carcinomas (N=11)** | 0%<br>(N=0) | 45.5%<br>(N=5) | 45.5%<br>(N=5) | 9%<br>(N=1) | 0%<br>(N=0) | 9%<br>(N=1) | 27.3%<br>(N=3) | 63.7%<br>(N=7) | 0%<br>(N=0) | 0%<br>(N=0) | 6.4 (± 2.7) |
| **Carcinomas Lymph Node Metastasis (N=10)** | 0%<br>(N=0) | 0%<br>(N=0) | 20%<br>(N=2) | 50%<br>(N=5) | 30%<br>(N=3) | 0%<br>(N=0) | 10% (N=1) | 40%<br>(N=4) | 40%<br>(N=4) | 10%<br>(N=1) | 10.1 (±2.2) |
| **Lymph Node Metastasis (N=10)** | 0%<br>(N=0) | 30%<br>(N=3) | 50%<br>(N=5) | 20%<br>(N=2) | 0%<br>(N=0) | 30%<br>(N=3) | 40% (N=4) | 30%<br>(N=3) | 0%<br>(N=0) | 0%<br>(N=0) | 11.2 (±2.0) |
| **Carcinomas Lung Metastasis (N=5)** | 0%<br>(N=0) | 0%<br>(N=0) | 0%<br>(N=0) | 60%<br>(N=3) | 40%<br>(N=2) | 0%<br>(N=0) | 0% (N=0) | 40%<br>(N=2) | 40%<br>(N=2) | 20%<br>(N=1) | 12.7 (±2.0) |
| **Lung Metastasis (N=5)** | 0%<br>(N=0) | 20%<br>(N=1) | 60%<br>(N=3) | 20%<br>(N=1) | 0%<br>(N=0) | 20%<br>(N=1) | 40% (N=2) | 0%<br>(N=0) | 20%<br>(N=1) | 20%<br>(N=1) | 11.3 (±1.7) |

\*SD: standard deviation

**Figure 3.**
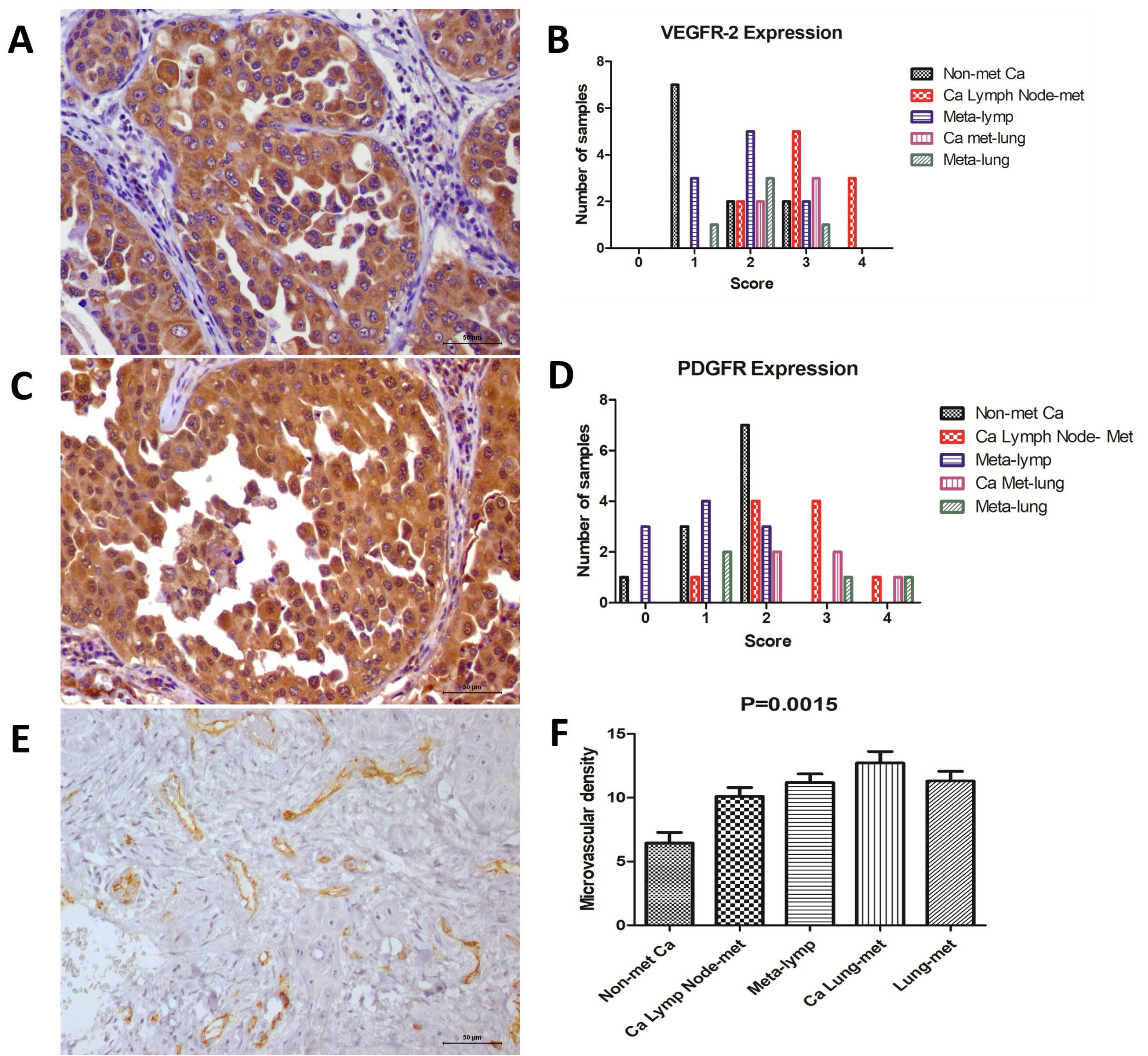
Immunoexpression of the different markers. A: VEGFR-2 (score 4) expression in a mammary carcinoma with lymph node metastasis. B: Graphic representation of each immunohistochemical score for VEGFR-2 expression in all tumor groups. C: PDGFR expression in a mammary carcinoma (score 2) with lymph node metastasis. D: Graphic representation of each immunohistochemical score for PDGFR expression in all tumor groups. E: Microvascular density (MVD) in a mammary carcinoma with lymph node metastasis. F: Graphic representation of MVD, indicating an increased number of vessels in metastatic tumors.

Regarding PDGFR immunoexpression, we observed positive expression in 25 out of 26 samples. No significant difference in PDGFR expression was noted among the different groups. No correlation was observed between PDGFR expression and tumor grade (P=0.0692; Spearman R=0.3620) or overall survival (P=0.2581; Spearman R= -0.2301). However, we identified a positive correlation between VEGFR and PDGFR expression. Thus, samples exhibiting the highest scores for VEGFR also presented the highest PDGFR expression (P=0.01; Spearman R=0.4959). PDGFR immunoexpression results are presented in Figure 3 and Table 2.

Regarding MVD (Figure 3), we identified an increased number of vessels in primary tumors with lymph node metastasis (P=0.0151) and lung metastasis (P=0.0046) compared with tumors with no metastasis. Primary tumors with lung metastasis exhibited increased MVD compared with carcinoma with lymph node metastasis (P=0.0496). Interestingly, we did not find a correlation between MVD and VEGFR expression (P=0.0827; Spearman R= 0.3467); however, a positive correlation between MVD and PDGFR was observed (P=0.0102; Spearman R= 0.4946) (Figure 4). Thus, samples with high PDGFR expression also exhibited high MVD (Figure 4).

**Figure 4.**
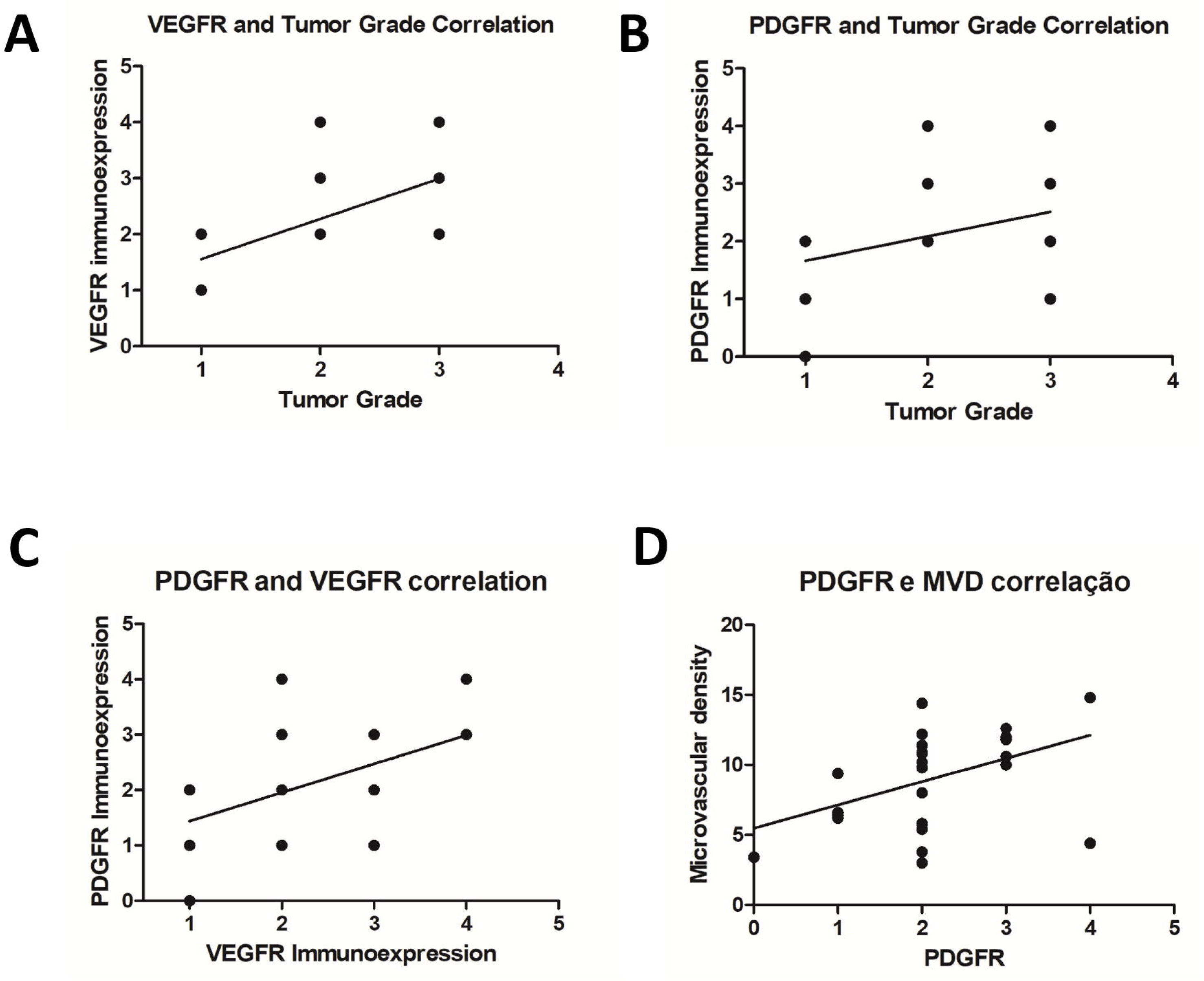
Correlation between immunohistochemical markers and different clinical parameters. A: Positive correlation between VEGFR-2 immunoexpression and tumor grade (P=0.001; Spearman R=0.6071). B: Absence of correlation between PDGFR expression and tumor grade (P=0.0692; Spearman R=0.3620). C: Positive correlation between VEGFR-2 and PDGFR expression (P=0.01; Spearman R=0.4959). D: Positive correlation between PDGFR expression and microvascular density (P=0.0102; Spearman R= 0.4946).

## 4. Discussion

Canine mammary gland tumors are one of the most important cancers in intact female dogs and represents a therapeutic challenge. Although surgery and chemotherapy have been used for CMT treatment, there is no standardized chemotherapy or target therapy. This research evaluated VEGFR-2, PDGFR2 and MVD in canine mammary tumors, aiming to associate different prognostic factors with these proteins. One interesting aspect of our research is a very restricted criterion used in patient selection. Typically, lymph node metastasis is evaluated after chain mastectomy in inguinal lymph nodes, and the inguinal lymph node is not always draining the tumor. The sentinel lymph node technique allowed us to identify the tumor-draining lymph node and increase the probability of identifying metastasis.

Another interesting aspect was the inclusion of a group of patients with no metastatic disease detected at diagnosis but with late lung metastasis. In clinical practice, it is relatively common to find female dogs with late lung metastasis after months or even years post surgery. However, given that metastatic disease takes a while to be detected on the X-ray, it was not possible to achieve a high number of patients in this group. These particular criteria can explain the highest survival time of patients with lung metastasis compared with patients with lymph node metastasis. We considered overall survival between the diagnosis and the time of current follow-up/death. Given that lung metastasis appeared in the patients from this group, the overall survival was compared with no lymph metastasis patients.

We did not include patients with more than one tumor in the mammary chain. This criterion excluded many animals from our study. Fifteen dogs were excluded from this study due to the presence of multiple mammary tumors (data not shown). In the context of multiple mammary tumors, it is not possible to guarantee which nodule the metastasis originated from. Regarding the nonmetastatic group, some patients with multiple tumors exhibited different molecular subtypes (data not shown), making it difficult to establish a prognosis based on the molecular subtype. Triple-negative tumors seem to be the most common molecular subtype in dogs [8]. This finding highlight the utility of dogs as a model for human triple-negative BC.

VEGFR-2 expression is correlated with angiogenesis and modulation of the tumor microenvironment [12]. In human [31] and canine [12] mammary tumors, VEGFR-2 expression is important in tumor growth and development and exhibits prognostic value [12]. Moreover, VEGFR-2 is a tyrosine kinase protein that can be inhibited by different target therapies [32]. Our results strongly suggested that VEGFR-2 is overexpressed in tumors with metastasis, indicating its predictive and prognostic value. Given that the VEGFR-2 inhibitor is not routinely used in human and veterinary oncology, clinical studies in dogs can benefit both species. We demonstrated a correlation between VEGFR-2 expression and tumor grade, indicating that high-grade tumors may require increased angiogenesis to maintain cell proliferation. Although we did not identify a correlation between VEGFR-2 expression and MVD, we identified increased vascular density in metastatic carcinomas.

These results together demonstrate the dependency of high-grade/metastatic tumors on angiogenic factors. Santos et al. [12] investigated VEGFR-2 expression in CMT, and overexpression of this protein was associated with carcinosarcomas (very aggressive tumor subtype). Although MVD did not correlate with VEGFR-2 expression, we identified a correlation between PDGFR and MVD. PDGFR induced intratumoral lymphogenesis [21], and we identified a correlation between intratumoral vasculogenesis and high levels of PDGFR. In addition, PDGFR and VEGFR-2 exhibited a positive correlation. These results collectively demonstrated the role of angiogenesis in the development and potential aggressiveness of CMT. Interestingly, both primary tumors and respective metastases were positive for VEGFR and PDGFR immunoexpression. Given that numerous VEGFR/PDGFR inhibitors are available, these results indicate the use of target therapy patients with CMT. Thus, our results support the idea of future clinical trials investigating the role of VEGFR/PDGFR inhibitors for the treatment of metastatic CMT.

## 5. Conclusions

Metastatic mammary carcinomas present VEGFR-2 overexpression and high microvascular density, indicating the role angiogenesis in tumor progression. PDGFR may induce vasculogenesis in metastatic mammary carcinomas. Overall, our results suggest the use of antiangiogenic and specific target therapies in a subset of patients with mammary tumors.

## Acknowledgments

We would like to thank the São Paulo Research Foundation (FAPESP) for their financial support (grant number: 2015/02798-5).

